# Stroke-associated pattern of gene expression previously identified by machine-learning is diagnostically robust in an independent patient population

**DOI:** 10.1101/109777

**Authors:** Grant C. O’Connell, Paul D. Chantler, Taura L. Barr

**Affiliations:** Center for Basic and Translational Stroke Research, Robert C. Byrd Health Sciences Center, West Virginia University, Morgantown, West Virginia; Department of Pharmaceutical Sciences, School of Pharmacy, West Virginia University, Morgantown, WV; Center for Cardiovascular and Respiratory Sciences, Robert C. Byrd Health Sciences Center, West Virginia University, Morgantown, West Virginia; Division of Exercise Physiology, School of Medicine, West Virginia University, Morgantown, West Virginia; Valtari Bio, Morgantown, WV

**Keywords:** ANTXR2, STK3, PDK4, CD163, MAL, GRAP, ID3, CTSZ, KIF1B, PLXDC2, GA/kNN

## Abstract

**Purpose:** Our group recently identified a ten gene pattern of differential expression in peripheral blood which may have utility for detection of stroke. The objective of this study was to assess the diagnostic capacity and temporal stability of this stroke-associated transcriptional signature in an independent patient population.

**Methods:** Publically available whole blood microarray data generated from 23 ischemic stroke patients at 3, 5, and 24 hours post-symptom onset, along with 23 cardiovascular disease controls were obtained via the National Center for Biotechnology Information Gene Expression Omnibus. Expression levels of the ten candidate genes were extracted, compared between groups, and evaluated for their discriminatory ability at each time point.

**Results:** We observed a largely identical pattern of differential expression between stroke patients and controls across the ten candidate genes as reported in our prior work. Furthermore, the coordinate expression levels of the ten candidate genes were able to discriminate between stroke patients and controls with levels of sensitivity and specificity upwards of 90% across all three time points.

**Conclusions:** These findings confirm the diagnostic robustness of the previously identified pattern of differential expression in independent patient population, and further suggest that it is temporally stable over the first 24 hours of stroke pathology.

## Introduction

The ability of clinicians to confidently recognize stroke during triage increases access to interventional treatments and affords patients improved odds for favorable outcome.^1, 2^ However, the diagnostic tools currently available to emergency medical technicians, paramedics, and hospital staff for identification of stroke have significant limitations.^3, 4^ Biomarker-based tests are clinically used to aid in the diagnosis of acute cardiovascular conditions such as myocardial infarction,^5^ however no such assay currently exists for the detection of stroke. This diagnostic limitation has resulted in a push for the identification of peripheral blood stroke biomarkers which could be rapidly measured in either the field or emergency department to guide early triage decisions.^3, 6^

Our group recently employed high-throughput transcriptomics in combination with a machine learning technique known as genetic algorithm/k-nearest neighbors (GA/kNN) to identify a panel of ten candidate genes whose peripheral blood expression levels were able to differentiate between 78 ischemic stroke patients and 74 control subjects with a high degree of accuracy.^7^ These candidate genes include seven whose expression levels were elevated in stroke patients relative to controls (*CD163, ANTXR2, PDK4, PLXDC2, STK3, ID3, CTSZ, KIF1B*), and three whose expression levels were down regulated (*MAL, ID3, GRAP*); their coordinate pattern of differential expression was able to discriminate between groups with levels of sensitivity and specificity approaching 100%. While the levels of diagnostic performance observed in this discovery investigation were unprecedented, limitations in study design necessitate further evaluation of the candidate genes in a validation analysis before definitive conclusions can be made regarding their true diagnostic efficacy.

Stroke patients and control subjects in this discovery investigation were not well matched in terms of cardiovascular disease (CVD) risk factors, leaving open the possibility that the pattern of differential expression which we observed across the ten candidate genes was driven by underlying CVD, and not by the acute event of stroke itself. Furthermore, subjects in this discovery study were almost exclusively Caucasian, and it is currently unknown whether ethnicity impacts the diagnostic efficacy the candidate genes, a possibility which deserves consideration due to the fact that there can be notable inter-ethnic differences in the pathophysiology of cardiovascular conditions.^8-11^ A further limitation in of this discovery study was the fact that blood samples were only collected at a single time point, making the temporal stability of candidate gene differential expression unclear with regards to the progression of stroke pathology. While post-hoc statistical analysis were used to address these potential confounds as best possible, it would be reassuring to observe similar levels of diagnostic performance across multiple time points in a more ethnically diverse subject pool which is better matched in terms of CVD risk factors.

Stamova *et al.* recently used microarray to examine gender differences in the response of the peripheral immune system to stroke.^12^ This investigation produced a publically available data set which includes genomewide whole blood expression data generated from 23 cardioembolic ischemic stroke patients at three replicate time points post-symptom onset (3, 5, and 24 hours), as well as from 23 neurologically asymptomatic control subjects; this patient population was ethnically diverse and groups were well matched in terms of risk factors for CVD. In the study reported here, we assessed the diagnostic robustness of the ten previously identified candidate genes in the aforementioned publicly available data set.

## Methods

### Data procurement

Raw whole blood microarray data (Affymetrix Human Genome U133 Plus 2.0 Array) were downloaded as CEL files from the National Center for Biotechnology Information (NCBI) Gene Expression Omnibus (GEO) via accession number 3000545. Patient demographic characteristics were aggregated from the gender-wise information reported by Stamova *et al.*^12^

### Microarray analysis

Analysis of microarray data was performed using the ‘affy’ package for R (R project for statistical computing).^13, 14^ Raw perfect match probe intensities were background corrected, quantile normalized (Figure 1), and summarized at the set level via robust multi-array averaging using the rma() function.^15^ Probe set level data associated with the ten candidate genes were then extracted for differential expression analysis; in the case of candidate genes with more than one associated probe set, data were further summarized at the gene level via simple averaging. Gene level summarized expression levels were then compared between stroke patients and controls across all three post-onset time points.

**FIGURE 1.**
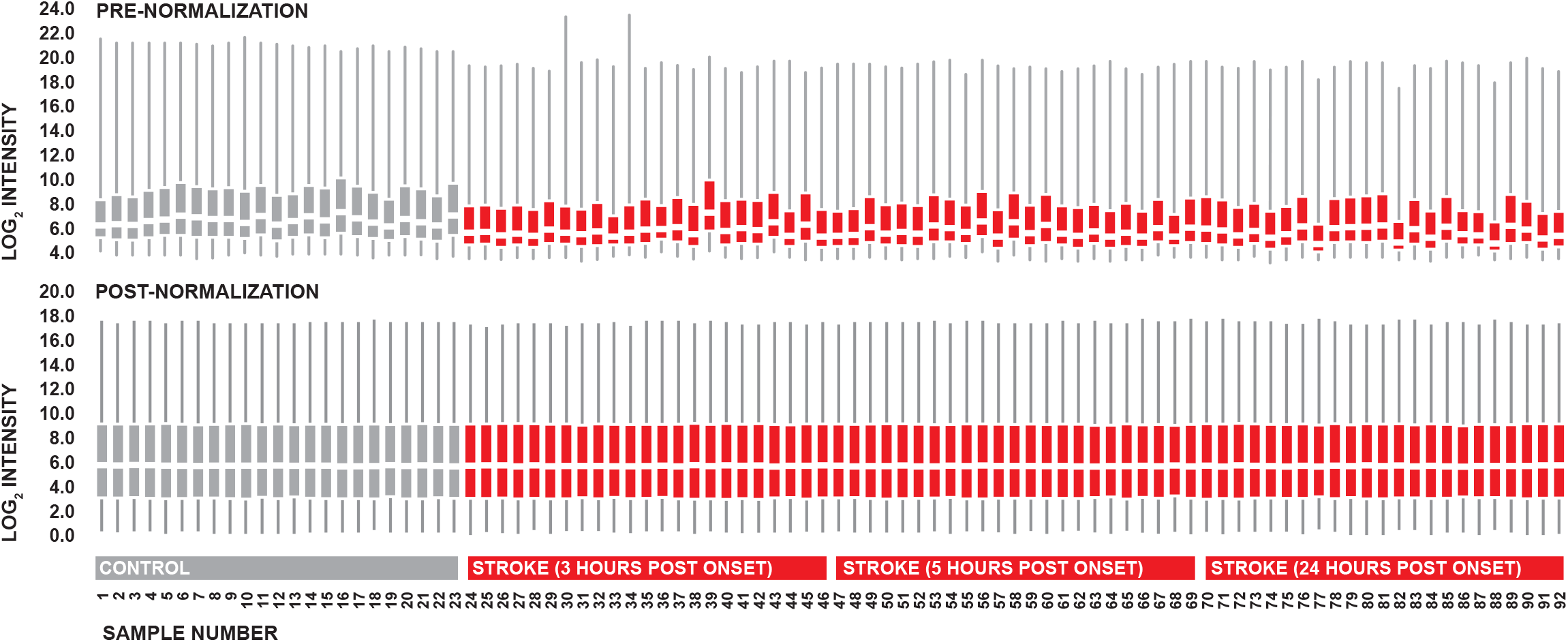
Normalization of microarray data. Distributions of pre and post-normalization perfect match probe intensities. Boxplots indicate standard five number summary values.

### Diagnostic evaluation

The diagnostic robustness of candidate gene expression levels was tested in terms of their ability to discriminate between stroke patients and controls using k-nearest neighbors (kNN) at each time point post-symptom onset. Classification was performed using standardized expression values, five nearest neighbors, and majority rule via the knn.cv() function of the ‘class’ package for R.^16^ The resultant prediction probabilities were used to generate receiver operator characteristic (ROC) curves using the roc() function of the ‘pROC’ package for R.^17^ Areas under the curves were then compared between time points via the roc.test() function according the non-parametric method described by DeLong *et* al.^18^

### Statistics

All statistics were performed using R 3.3. Fisher’s exact test was used for comparison of dichotomous variables. T-test or one-way ANOVA was used for comparisons of continuous variables where appropriate. The null hypothesis was rejected when p<0.05. In the case of multiple comparisons, p-values were adjusted via Holm’s Bonferroni correction.^19^

## Results

### Clinical and demographic characteristics

Stroke patients were significantly older than control patients, but well matched in terms of gender and ethnicity. In terms of cardiovascular disease risk factors, groups were well matched with regards to rates of hypertension and diabetes, however control subjects displayed a significantly higher prevalence of dyslipidemia relative to stroke patients (Table 1).

**TABLE 1.**
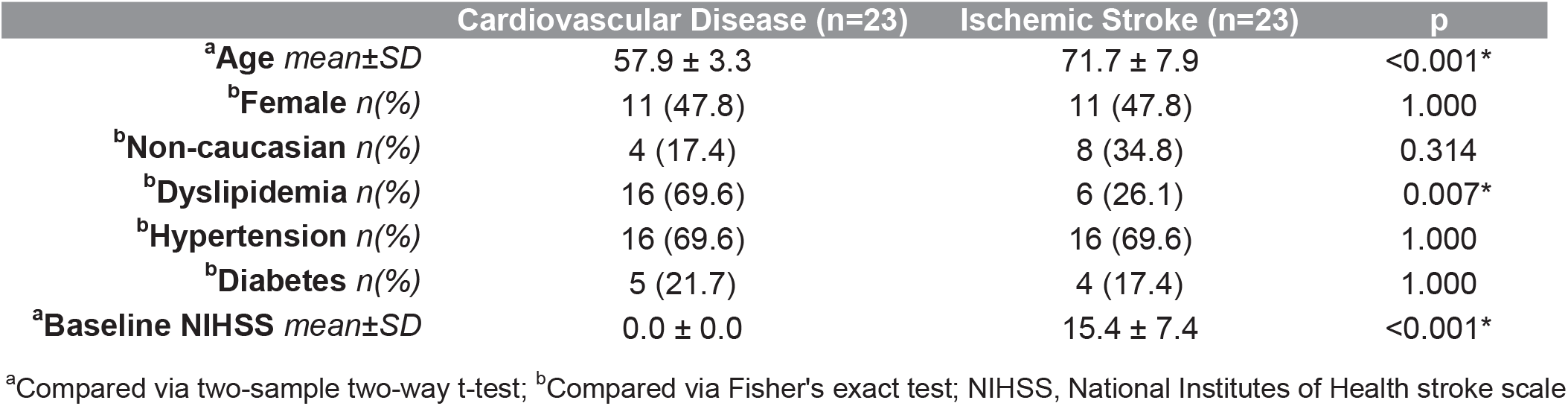
Clinical and demographic characteristics.

### Microarray data processing

Distributions of perfect match probe intensities were visually similar following normalization, providing indication that normalized expression data were suitable for intersample comparison (Figure 1). Probe sets extracted for differential expression analysis are listed in Table 2.

**TABLE 2.** Probe sets extracted for differential expression analysis.

| Gene | Affy Probe Set ID | Target Transcripts<sup>a</sup> | Gene | Affy Probe Set ID | Target Transcripts<sup>a</sup> |
| --- | --- | --- | --- | --- | --- |
| **CD163** | 203645_s_at | NM_004244 | **STK3** | 204068_at | NM_001256312 |
|  | 215049_x_at | NM_203416 |  | 211078_s_at | NM_001256313 |
|  | 216233_at | XM_005253528 |  |  | NM_006281 |
|  |  | XM_005253529 |  |  | XM_005251034 |
|  |  | XR_429039 |  |  |  |
| **ANTXR2** | 1555536_at | NM_001145794 | **CTSZ** | 210042_s_at | NM_001336 |
|  | 225524_at | NM_001286780 |  | 212562_s_at |  |
|  | 228573_at | NM_001286781 | **GRAP** | 206620_at | NM_006613 |
|  | 238050_at | NM_058172 |  | 229726_at | XM_005256425 |
| **PDK4** | 205960_at | NM_002612 |  |  | XM_005256426 |
| **PLXDC2** | 214807_at | NM_001282736 | **KIF1B** | 209234_at | NM_015074 |
|  | 226865_at | NM_032812 |  | 225878_at | NM_183416 |
|  | 227276_at |  |  | 226968_at |  |
|  | 227995_at |  |  | 228657_at |  |
|  | 236297_at |  | **MAL** | 204777_s_at | NM_002371 |
|  | 238455_at |  |  |  | NM_022438 |
| **ID3** | 207826_s_at | NM_002167 |  |  | NM_022439 |
|  |  |  |  |  | NM_022440 |

<sup>a</sup>NCBI RefSeq ID

### Candidate gene differential expression

Six of the seven candidate genes which we had previously reported as being elevated in stroke in our prior investigation displayed similar up-regulation in stroke patients relative to controls (Figure 2A, 2B, 2D, 2E, 2F, 2J), however one exhibited no significant differences in expression levels at any time point post-symptom onset (Figure 2H). In terms of the candidate genes which we had previously reported as being down regulated in stroke, all three displayed significantly lower expression levels in stroke patients relative to controls (Figure 2C, 2G, 2I). Collectively, these observations largely confirmed the pattern of candidate gene differential expression reported in our prior investigation.

**FIGURE 2.**
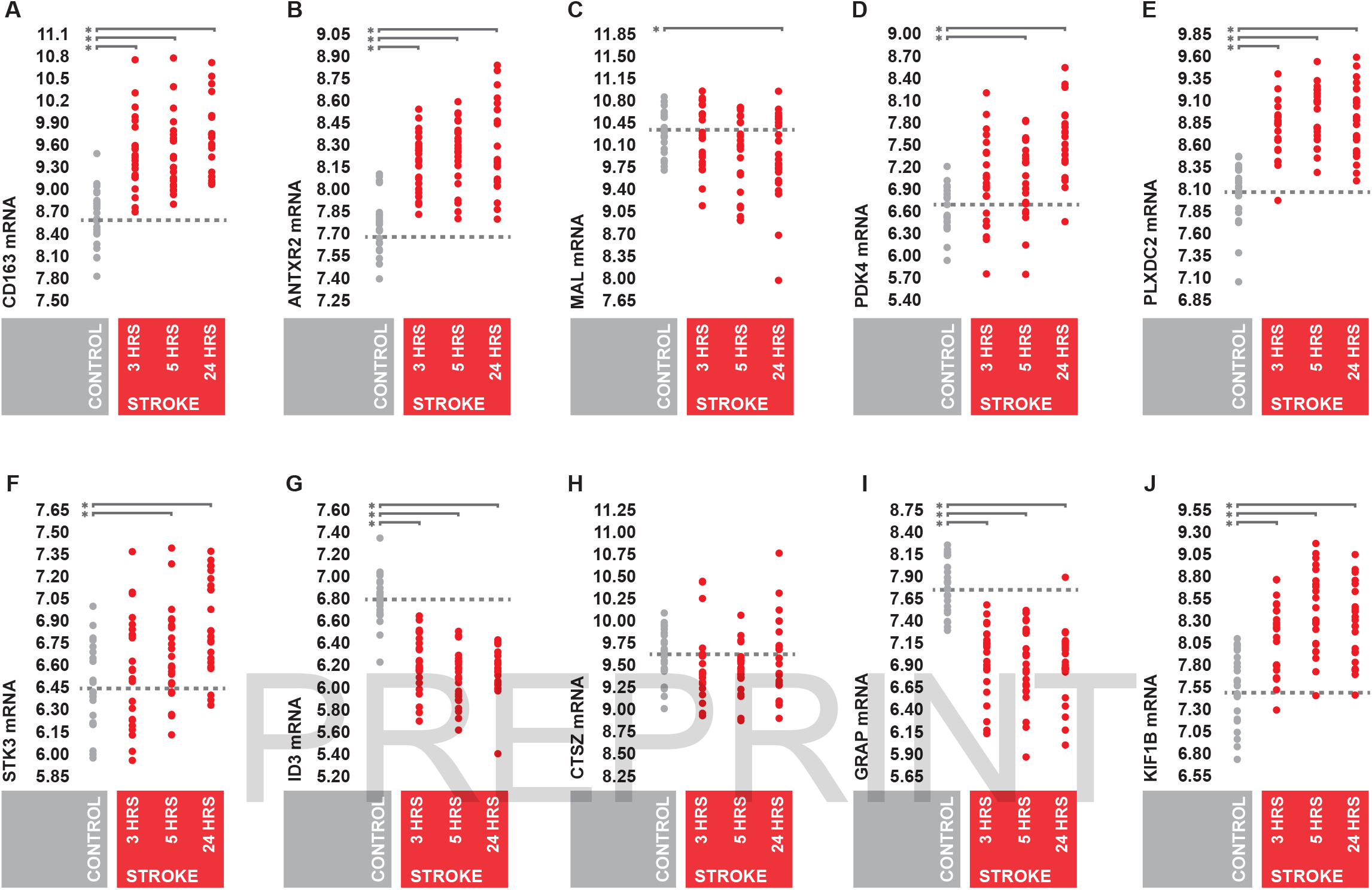
Candidate gene differential expression. (A-J) Peripheral whole blood expression levels of candidate genes in stroke patients and controls at 3, 5, and 24 hours post-symptom onset. Expression values represent gene-level summarized log_2_ perfect match probe intensities. Expression levels were statistically compared between stroke patients and controls across all time points using one-way AVOVA. Post-hoc testing was performed via two-sample two-way t-test with Holm’s Bonferroni correction for multiple comparisons.

### Temporal profile of candidate differential expression

Most candidate genes displayed some degree of differential expression by three hours post-symptom onset, and the magnitude of the overall response appeared to increase over time. Several candidate genes appeared to achieve maximal differential expression at five hours post-onset and then plateau, while a few displayed steady increases in the degree of differential expression through 24 hours (Figure 3), providing evidence that the expression levels of the candidate genes are likely directly responsive to acute stroke pathology.

**FIGURE 3.**
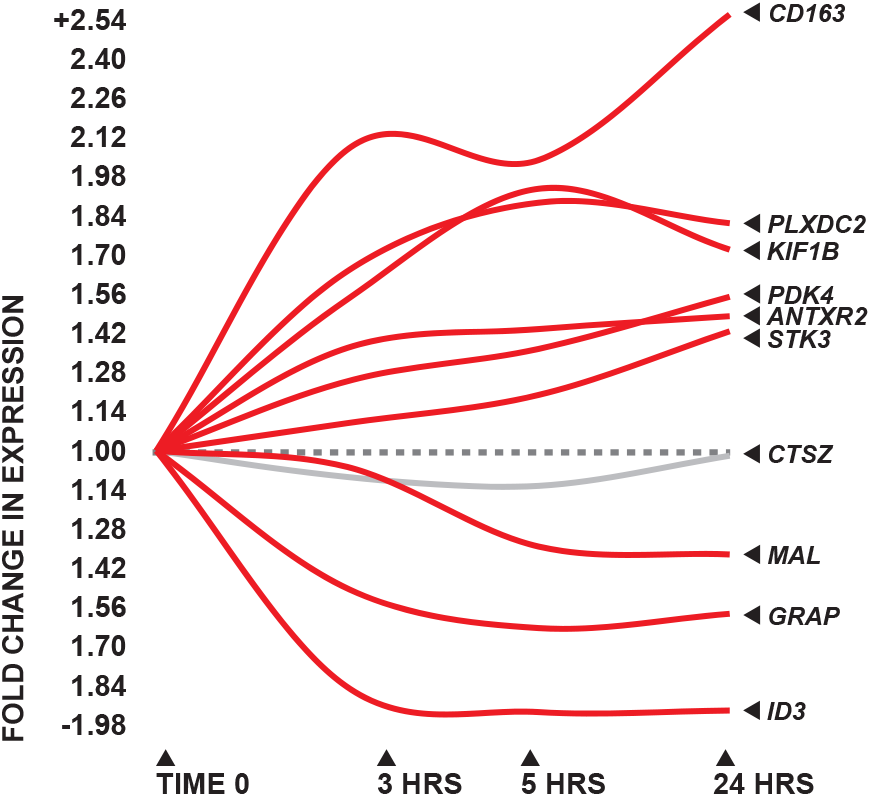
Temporal profile of candidate gene differential expression. Magnitude of candidate gene differential expression between stroke patients and controls at 3, 5, and 24 hours post-symptom onset, indicated as fold difference relative to control.

### Candidate gene diagnostic performance

In terms of diagnostic ability, the coordinate expression levels of the ten candidate genes were able to discriminate between stroke patients and controls using kNN with levels of sensitivity and specificity upwards of 90% at all three time points post-symptom onset (Figure 4A, 4B, 4C). While the overall diagnostic capacity of the ten candidate genes appeared slightly more robust at five and 24 hours, no statistically significant differences in area under ROC curve were observed between time points (Figure 4D). Taken together, these observations supported the high levels of diagnostic performance reported in our prior work, and suggest that the diagnostic capacity of the ten candidate genes is temporally stable over the first 24 hours post-symptom onset.

**FIGURE 4.**
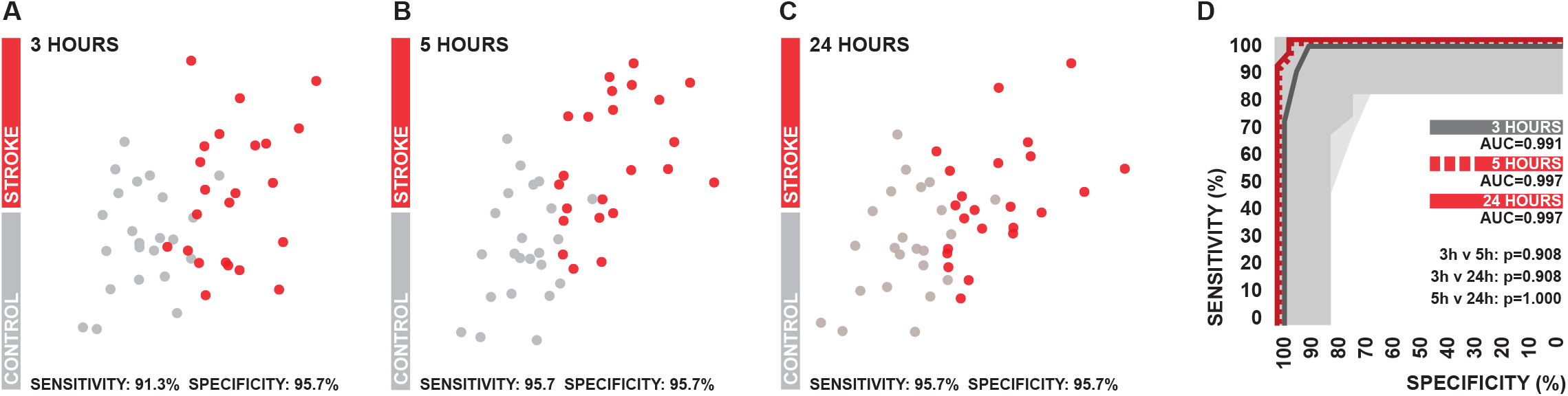
Candidate gene diagnostic performance. (A-C) Two-dimensional projections of the kNN feature spaces generated by the coordinate expression levels of the ten candidate genes at 3, 5, and 24 hours post-symptom onset. Levels of sensitivity and specificity are associated with class predications generated via five nearest neighbors using a probability cutoff of 0.50. (D) ROC curves associated with the prediction probabilities generated in kNN. Shaded areas indicate 95% confidence intervals. Areas under curves were statistically compared between time points using the DeLong method and p-values were adjusted using Holm’s Bonferroni correction to account for multiple comparisons.

## Discussion

There has been a recent push for the identification of circulating biomarkers which could be used to aid clinicians in the recognition of stroke during patient triage. Our group recently employed high-throughput transcriptomics in combination with a machine learning technique known at GA/kNN to identify a ten gene pattern of differential expression in peripheral blood which has potential utility for the detection of stroke.^7^ However, patients in this discovery investigation were almost exclusively Caucasian, groups were not well matched in terms of CVD risk factors, and blood was only sampled at a single time point post-symptom onset. In the study reported here, we leveraged a publically available microarray dataset to evaluate the previously identified candidate pattern of gene expression at multiple pathological time points in a more ethnically diverse subject pool which was better matched in terms of CVD risk factors.

The overall pattern of differential expression which we previously reported between stroke patients and controls was largely confirmed in the analysis described here, as nine of the ten candidate genes were identically differentially regulated. Furthermore, the candidate genes displayed similar levels of diagnostic robustness as described previously. This suggests that it is unlikely that our prior findings were substantially driven by intergroup differences in CVD risk factors; this notion is accentuated by the fact that the overall pattern of differential expression across the ten candidate genes was temporally dynamic with regards to time from symptom onset, providing evidence that the candidate genes are directly responsive to stroke pathology. The fact that our prior observations were largely recapitulated in the analysis reported here also suggests that ethnicity likely has little influence on the overall transcriptional response of the candidate genes to stroke.

One possible exception with this regard is *CTSZ*, which was the only candidate gene which failed to exhibit a similar response to stroke as previously reported. Thus, it is possible that the differential regulation of *CTSZ* which we observed in our discovery investigation was indeed driven primarily by underlying CVD, or that there are interethnic differences in the responsiveness of *CTSZ* to stroke. However, to our knowledge, there are no associations reported in the literature to support either conclusion, and is possible that the discrepancy in response between investigations has other explanation. The samples analyzed in this study were obtained exclusively from patients presenting with ischemic strokes of cardioembolic etiology, while the samples used in our prior discovery study were obtained from patients presenting with ischemic strokes of multiple etiologies, including a large number which were thrombotic in nature; thus it is possible that the disagreement in findings is due to an etiology-specific response. The disagreement in findings could also be driven by a technical confound, as the gene expression data used in this analysis were generated using a different gene chip then that which was used in our discovery investigation, and the chips do not have completely overlapping transcriptional coverage of *CTSZ*.

In addition to providing a general validation of the overall pattern of candidate gene differential expression, this study also afforded us an opportunity to evaluate its temporal stability with regards to stroke pathophysiology. The overall pattern of differential expression was modestly detectable at three hours post-symptom onset and appeared to increase in magnitude though 24 hours. Despite the modest magnitude, the levels of differential expression present at three hours post-onset were still adequate to differentiate between groups with similarly high levels of diagnostic performance as those observed at the subsequent two time points. Overall, our findings suggested that the diagnostic ability of the candidate pattern of gene expression is relatively temporally stable over the first 24 hours of stroke pathophysiology, which is encouraging from a translational standpoint in that the first clinical contact with stroke patients tends to vary across a wide time range with regards to time from onset, depending in the overtness of symptom presentation.

A potential limitation with regards to this study lies in that the stroke patients and controls associated with the samples used in this analysis were not well matched in terms of age. Ideally, multiple regression could be used to statistically control for such a potential confound, however non-aggregated demographic information was not available for the dataset, making such an analysis impossible. However, we explored the relationship between the expression levels of the ten candidate genes and age as part of our previously reported discovery investigation, and observed no significant associations. Thus, we feel that it is unlikely that the results reported here are confounded by intergroup age differences.

Collectively, the findings of this analysis confirm the diagnostic robustness of the previously identified stroke-associated pattern of gene expression, and further suggest that it is temporally stable over the first 24 hours of stroke pathology. Due to fact that this transcriptional signature has now demonstrated levels of diagnostic performance which well exceed those of the triage tools currently available to clinicians for identification of stroke in two independent investigations, we feel that it has legitimate translational potential and a path towards clinical implementation should be further explored.

## Contributions

Work was conceptualized by GCO. Analysis was performed by GCO. Work was financially supported by TLB and PDC. Manuscript was written by GCO with contributions from TLB and PDC.

